# A Conserved Stress-Inducible Hub Enhancer Governs Natriuretic Peptide Expression and Undergoes Dynamic and Reversible Activation in Human Heart Failure

**DOI:** 10.1101/2025.10.13.682224

**Authors:** Ken Matsuoka, Hijiri Inoue, Hyejin Jo, Chisato Okamoto, Hirona Hasuike, Hai Ying Fu, Hidetaka Kioka, Yuki Kuramoto, Takatsugu Segawa, Shunsuke Nishimura, Rina Takamiya, Yohei Miyashita, Yoshihiro Asano, Satoru Yamazaki, Hisakazu Kato, Yasushi Sakata, Seiji Takashima, Osamu Tsukamoto

## Abstract

**Background:** Natriuretic peptides (NPs), encoded by the *NPPA* and *NPPB* genes, serve as both diagnostic biomarkers and cardioprotective hormones in heart failure. Their expression is tightly regulated by mechanical load, yet the upstream enhancer mechanisms translating hemodynamic stress into transcriptional activation remain incompletely understood.

**Methods:** We investigated the *Nppa*/*Nppb* super-enhancer (SE), which resides in a well-insulated topologically associating domain (TAD) and regulates stress-inducible expression of *Nppa* and *Nppb*. We applied CRISPR-based enhancer perturbation, chromatin accessibility and histone profiling, chromatin conformation assays, genetic mouse models, human iPSC-derived cardiomyocytes, and paired failing human hearts before and after left ventricular assist device (LVAD) unloading.

**Results:** Through integrative epigenomic and genetic analyses, we identified a conserved element, CR9, as a dominant hub enhancer within this SE. In neonatal rat cardiomyocytes, CRISPR-based activation and inhibition established that CR9 was both necessary and sufficient for NP induction, whereas neighboring elements (CR7/CR8) displayed modest activity but synergized with CR9, establishing a hierarchical and cooperative SE architecture. In vivo, CR9 deletion in mice markedly suppressed *Nppa*/*Nppb* expression and diminished chromatin accessibility and histone acetylation across adjacent enhancers and promoters, underscoring its structural as well as transcriptional role. Reinsertion of CR9, even in reverse orientation, restored transcription, confirming its orientation-independent activity. CR9 function remained confined within TAD boundaries, supporting a locus-specific regulation. RNAscope revealed spatial proximity between CR9 enhancer RNA and *Nppa*/*Nppb* nascent transcripts. Developmental profiling revealed a switch in enhancer usage from CR6/CR7 in the fetal heart to CR9 in adulthood, indicating a transition from developmental to stress-inducible regulation. In human induced pluripotent stem cell–derived cardiomyocytes, deletion of CR9 nearly abolished *NPPA*/*NPPB* expression, establishing its essential role in a human cellular context. Most notably, in paired human failing hearts before and after LVAD unloading, chromatin accessibility at CR9 was dynamically reversed, providing the first direct evidence that enhancer states are plastic and therapeutically modifiable in the human myocardium.

**Conclusions:** CR9 functions as a stress-inducible hub enhancer that coordinates NP transcription under pathological load. This enhancer exhibits reversible activation in human hearts, underscoring enhancer plasticity as a modifiable regulatory layer mediating stress-responsive transcriptional adaptation in the diseased myocardium.

**Clinical Perspective:** *What is new?:*

- We identified CR9 as a stress-inducible hub enhancer within the *Nppa*/*Nppb* super-enhancer that is both necessary and sufficient for natriuretic peptide induction, defining a hierarchical and cooperative regulatory architecture controlling *Nppa* and *Nppb* expression.
- We uncovered a developmental switch in enhancer usage from CR6 in the fetal heart to CR9 in adulthood, revealing how stress-inducible enhancer activity emerges as the heart transitions from developmental to adaptive regulation.
- Using paired failing human hearts before and after LVAD unloading, we provide the first direct evidence that enhancer states are dynamically reversible in the human myocardium.

*What are the clinical implications?:*

- The reversibility of CR9 activity in human heart failure provides a molecular basis for how natriuretic peptide levels decrease when cardiac function improves with effective therapy, reflecting the underlying enhancer plasticity of the diseasesd myocardium.
- Enhancer plasticity is an emerging and targetable mechanism in heart failure, raising the possibility that stress-inducible enhancers like CR9 could be engineered to switch on cardioprotective genes when the heart is under stress, offering a new strategy for precision gene therapy.

## Introduction

Circulating natriuretic peptides, including atrial natriuretic peptide (ANP, encoded by *NPPA*) and B-type natriuretic peptide (BNP, encoded by *NPPB*), are indispensable biomarkers for the diagnosis, risk stratification, and therapeutic monitoring of heart failure, as endorsed by the 2022 AHA/ACC/HFSA Guideline for the Management of Heart Failure^1^. The *NPPA* and *NPPB* expression is minimal under physiological conditions but is robustly reactivated upon mechanical or neurohumoral stress in proportion to the severity of cardiac dysfunction^2–4^. Plasma ANP/BNP levels rise in decompensated heart failure and decline with effective therapy, consistent with their routine use as clinical surrogates of hemodynamic stress. Notably, in end-stage heart failure, myocardial *NPPA*/*NPPB* expression can be fully normalized by mechanical unloading via a left ventricular assist device (LVAD), underscoring their transcriptional plasticity to hemodynamic forces^5^ and providing a unique human model to study stress-inducible transcriptional regulation in vivo. These features make *NPPA*/*NPPB* ideal genes for studying stress-inducible transcriptional control in the heart. However, despite the clinical centrality of these peptides, the molecular mechanisms by which cardiomyocytes selectively and robustly induced *NPPA*/*NPPB* transcription under pathological stress remain poorly understood.

Enhancers are key regulatory DNA elements that drive cell type– and condition-specific gene expression^6–8^. Super-enhancers (SEs) represent large clusters of enhancers enriched for transcription factors, mediators, and active histone markers such as H3K27ac, and are thought to regulate cell identity and disease-relevant genes^8–11^. Although SEs are often treated as unified regulatory units, functional dissection studies have revealed heterogenous internal logic, ranging from additive^12,13^, redundant^14^, synergistic^15^, and hierarchical behaviors^16,17^. Importantly, *NPPA* and *NPPB* themselves display striking cardiac specificity and robust stress inducibility, features that strongly implicate enhancer-mediated regulation in their transcriptional control.

We previously identified a ∼650-bp conserved element, termed conserved region 9 (CR9), located ∼22 kb upstream of *Nppb*, as a conserved candidate enhancer robustly activated by mechanical stress in cardiomyocytes^2,3^. Subsequent study reported that CR9 resides within a larger ∼12-kb conserved enhancer cluster, designated regulatory element 1 (RE1), which has been classified as a cardiac -specific SE active during both development and pathological stress^4^. While stress-induced activation of CR9 parallels *Nppa*/*Nppb* induction, prior works have not established a causal relationship^2,3^. Moreover, RE1 contains additional conserved elements such as CR6, CR7 and CR8,^4^ but how these candidate enhancers functionally interact with CR9—whether they act redundantly, synergistically, or hierarchically—remains unknown in the native chromatin context, particularly in vivo. As highlighted in recent discussions on SE biology^8^, profiling approaches such as ChIP-seq are insufficient to predict functional output, underscoring the need for element-by-element dissection in vivo.

The *NPPA*/*NPPB* locus resides within a well-insulated topologically associating domain (TAD) and contains a cardiac-specific SE, along with two adjacent stress-inducible target genes, *NPPA* and *NPPB*. This configuration provides a well-defined framework to investigate the regulatory logic and internal architecture of a stress-responsive SE under physiological and pathological conditions. In this study, we set out to dissect the causal role of CR9 in natriuretic peptide gene induction and to determine how it cooperates with neighboring enhancer elements within the SE. By integrating CRISPR-based enhancer perturbation, chromatin accessibility and histone profiling, chromatin conformation analyses, genetic mouse models, human iPSC-derived cardiomyocytes, and paired human failing hearts before and after LVAD unloading, we demonstrate that CR9 functions as a stress-inducible hub enhancer that orchestrates cooperative regulation of *NPPA/NPPB* and exerts a dominant regulatory influence within the enhancer network under cardiac stress, with direct relevance to human heart failure and highlighting enhancer plasticity as a potential therapeutic axis.

## Methods

### Study approval

All animal experiments were performed following the Guide for the Care and Use of Laboratory Animals (NIH Publication, 8th Edition, 2011) and were approved by the Animal Care and Use Committee of the Osaka University Graduate School of Medicine. Testing of human samples was approved by the Ethics Committee of Osaka University Hospital (10081(T1)-27), and written informed consent was obtained from all patients before inclusion in the study.

### Experimental Models

Neonatal rat ventricular myocytes, genetically modified mice, human iPSC-derived cardiomyocytes (iPSC-CMs), and paired failing human hearts before and after LVAD unloading were analyzed. Primary neonatal rat ventricular myocytes were isolated and cultured under standard conditions, as previously described^9^, and detailed in Supplementary Methods. Animal experiments were performed at the Institute of Experimental Animal Sciences, Faculty of Medicine, The University of Osaka. For animal studies, sample sizes were based on previous similar studies. Each mouse represented one biological replicate. Animals were excluded only in cases of technical failure. Randomization was not applicable because experimental groups were defined by genotype. Blinding was not applied to experimental procedures; however, quantitative imaging and RNA analyses were performed objectively. The outcome measures were *Nppa*/*Nppb* expression, chromatin accessibility, and histone modifications.

### CRISPR-based enhancer perturbation

CRISPR activation (CRISPRa) and interference (CRISPRi) experiments were conducted to define enhancer function in cultured neonatal rat cardiomyocytes using previously established systems^18^. CRISPRa was achieved using dCas9–VP64 and CRISPRi using dCas9-KRAB or dCas9-LSD1 systems. Viral delivery and sgRNA targeting were performed as described in Supplementary Methods (Supplementary Figure 2). Detailed sgRNA sequences, cloning strategy, and vector information are listed in Supplementary Table 1.

### Generation of CR9 knockout and knock-in models

Human CR9 knockout iPSC line was generated as described in Supplementary Methods. CR9 knockout (CR9⁻/⁻) mice were produced via homologous recombination, and CR9 knock-in lines restoring the 650 bp CR9 sequence were generated using single-stranded oligodeoxynucleotides (ssODNs) and CRISPR/Cas9. Both forward (CR9^Fw/Fw^) and reverse (CR9^Rv/Rv^) orientations were obtained. Full genotyping and validation procedures are provided in Supplementary Methods (Supplementary Figure 5–6).

### Chromatin and transcriptomic profiling

Cardiomyocyte nuclei were isolated from frozen LV tissue by PCM-1-based FACS sorting^4^. Chromatin accessibility, histone modifications, and chromatin architecture were profiled by Assay for transposase-accessible chromatin with sequencing (ATAC-seq), CUT&Tag (H3K27ac and H3K4me3), and circular chromosome conformation capture (4C-seq), respectively. RNA-seq was performed on LV tissue, and differentially expressed genes (DEGs) were defined using an FDR < 0.05. Further details are provided in the Supplementary Methods.

### RNAscope fluorescence in situ hybridization (FISH) and imaging analysis

RNAscope fluorescence in situ hybridization (FISH) was performed on cultured rat cardiomyocytes to visualize nascent transcripts of *Nppa, Nppb, Myh6*, and CR9-derived eRNA using intronic probes. Confocal imaging and spatial quantification were performed using established methods^19^. RNAscope images were acquired using a confocal laser scanning microscope (Olympus FV-3000), and the spatial proximity between CR9 eRNA and *Nppb* nascent transcripts was quantified by calculating the 2D distances between signal centroids, as described previously^20^. Full imaging and quantification procedures are detailed in the Supplementary Methods.

### Human heart sample collection

LV tissues were collected from dilated cardiomyopathy patients who underwent LVAD surgery. Chromatin accessibility was analyzed using ATAC-seq. RNA expression was analyzed using qRT-PCR.

### Statistical analysis

All data are expressed as mean ± standard deviation (s.d.) unless otherwise indicated. For comparisons between two groups, a two-tailed Student’s *t*-test or Mann-Whitney *U* test was used. For comparisons among multiple groups, one-way analysis of variance (ANOVA) using Dunnett’s or Tukey’s comparison test was applied. Two-way analysis of variance (ANOVA) with Tukey’s post hoc test was used for murine experiments with or without PE treatment. The correlation between CR9 eRNA intensity and *Nppb* mRNA intensity was assessed using Pearson’s correlation coefficient. A *P* < 0.05 was considered statistically significant. Statistical analyses and graph generation were performed using the GraphPad Prism software (MDF; Tokyo, Japan).

### Data Availability

The sequencing data generated in this study will be deposited in the GEO database prior to publication in Supplementary Table 2. All the other data are available from the corresponding author upon request.

## Results

### CR9 functions as a dominant enhancer of *Nppa* and *Nppb* in cardiomyocytes

The *Nppa*–*Nppb* genomic cluster is located within a ∼47-kbp region flanked by two CTCF binding sites and lies within a single TAD that also contains adjacent genes, such as *Miip*, *Mfn2*, and *Plod1*^4^ (Figure 1A, Supplementary Figure 1A). We had previously identified CR9 as a cardiac-specific, stress-responsive putative enhancer among nine conserved noncoding elements (CR1–CR9)^2,3^. Notably, CR7 and CR8 are also located within RE1, which was previously characterized as a SE regulating *Nppa* and *Nppb*^4^. Similar to CR9, both elements exhibited distinct H3K27ac peaks, suggesting an active chromatin enhancer state.

**Figure 1.**
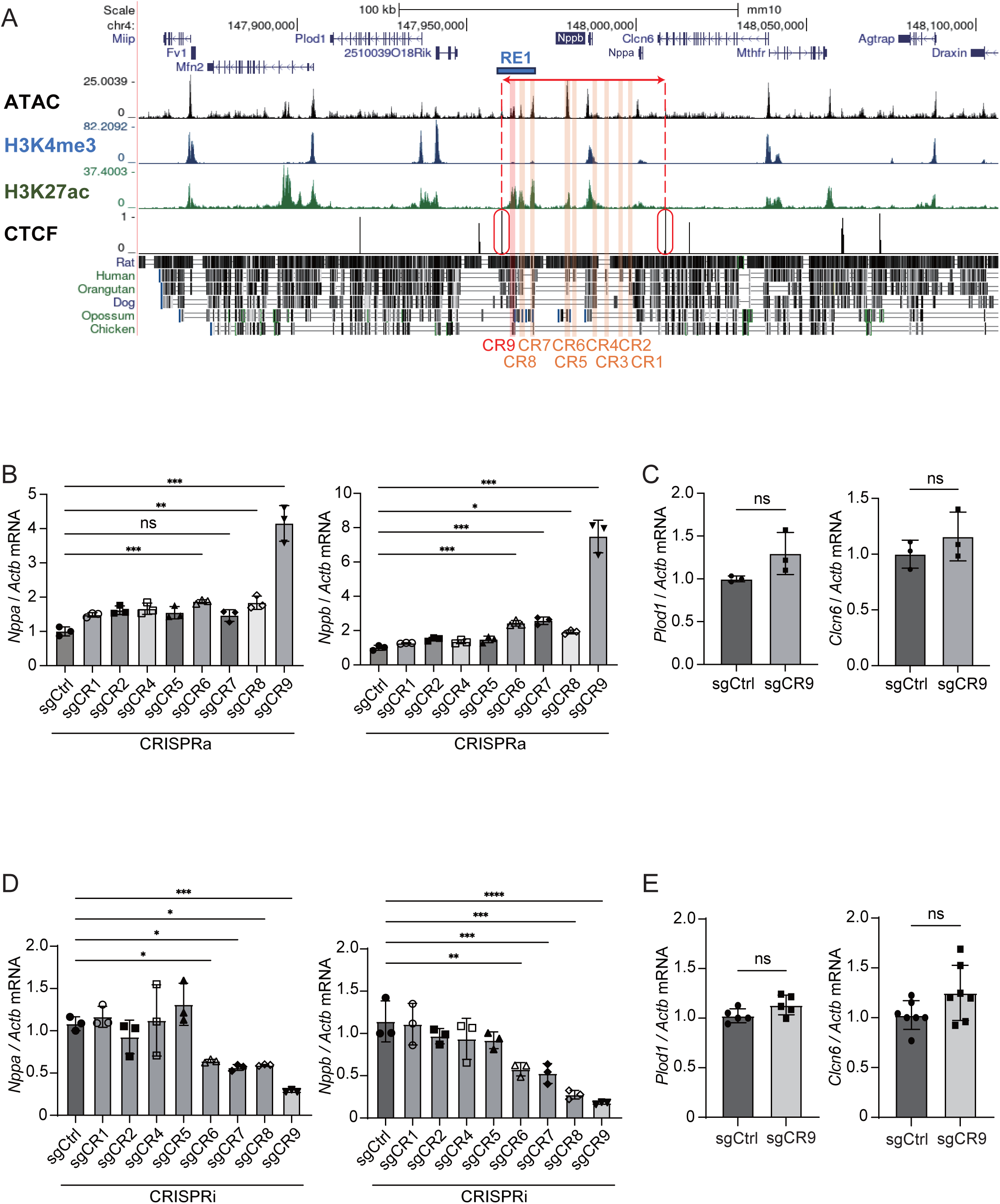
CRISPR-based epigenetic editing identifies CR9 as a dominant enhancer of *Nppa* and *Nppb* in neonatal rat cardiomyocytes. **(A)** UCSC Genome Browser view of the mouse *Nppa-Nppb* loci showing ATAC-seq, H3K4me3 ChIP, H3K27ac ChIP, and CTCF ChIP-seq data from adult murine hearts. RE1 enhancer cluster is highlighted in blue. Noncoding conserved regions (CR1–CR9), each at least 1 kb away from the transcription start sites of *Nppa* and *Nppb* and conserved between mouse and human, are shown in red to orange. **(B)** qPCR analysis of *Nppa* and *Nppb* mRNA expression in neonatal rat cardiomyocytes transduced with CRISPR activation (CRISPRa) constructs targeting CR1 to CR9 (sgCR1–sgCR9) or a control sgRNA (sgCtrl). Expression levels were normalized to *Actb* and shown as fold change relative to sgCtrl. **(C)** Expression of adjacent genes *PlodI* and *Clcn6* following CR9 activation by CRISPRa. **(D)** Expression of *Nppa* and *Nppb* following CRISPR interference (CRISPRi) targeting of CR1 to CR9. **(E)** Expression of *Plod1* and *Clcn6* following CRISPRi targeting of CR9. Data represent mean ± s.d.; *n* = 3 biologically independent experiments for (B–D); *n* = 5 (Plod1) and *n* = 7 (Clcn6) biologically independent samples for (E). Statistical significance was determined by one-way ANOVA with Tukey’s post hoc test for (B, D) and by two-tailed unpaired Student’s t-test for (C, E). \**P* < 0.05; \*\**P* < 0.01; \*\*\**P* < 0.001; \*\*\*\**P* < 0.0001; ns, not significant.

To define the causal role of CR9 in the transcriptional activation of *Nppa* and *Nppb*, we first employed CRISPR-based epigenetic perturbation strategies^18^ in cultured neonatal rat cardiomyocytes. Given that CR9 has been previously identified as a stress-inducible element embedded within the RE1 super-enhancer region^2–4^, we systematically compared its endogenous enhancer activity to that of neighboring elements (CR1–CR8, especially CR7 and CR8). To assess the regulatory contribution of each element, we individually targeted the CR elements using CRISPR activation (CRISPRa) with dCas9-VP64 and sequence-specific single-guide RNAs (sgRNAs)^18^ (Supplementary Figure 2A). Among the tested elements, only CR9 activation led to a robust upregulation of both *Nppa* and *Nppb* mRNA levels, whereas CR6, CR7 and CR8 had minimal or no effect (Figure 1B). Importantly, the expression of nearby genes, such as *Plod1* and *Clcn6*, remained unchanged, indicating the specificity of the CR9 enhancer (Figure 1C).

Next, we performed CRISPR interference (CRISPRi) using a dual-repressor CRISPRi system (dCas9-KRAB-LSD1) to assess the requirement of each enhancer element^18^ (Supplementary Figure 2B). Inhibition of CR9 led to a 70–80% reduction in *Nppa* and *Nppb* expression, whereas inhibition of CR6, CR7 and CR8 resulted in a more modest 50–60% reduction (Figure 1D). Targeting the other CR elements had no significant effect (Figure 1D). Similar to the CRISPRa results, CRISPRi of CR9 did not alter the expression of the adjacent genes, *Plod1* and *Clcn6* (Figure 1E), further supporting the element-specific regulatory role of CR9 in controlling *Nppa* and *Nppb* expression.

### CRISPRa/i of CR9 modulates enhancer–promoter chromatin states at the *Nppa–Nppb* locus

To investigate how CRISPR-based modulation of CR9 affects the epigenetic landscape at the *Nppa–Nppb* locus, we performed ChIP-seq for H3K27ac and H3K4me3, which mark active enhancers and promoters, respectively^21^. CRISPRa targeting of the CR9 region led to an increase in H3K27ac signal across the CR7–CR9 region, with no notable changes at non-targeted genomic sites (Figure 2A). Importantly, H3K4me3 and H3K27ac levels at the promoters of *Nppa* and *Nppb* were elevated following CR9 activation, consistent with transcriptional upregulation (Figure 2A).

**Figure 2.**
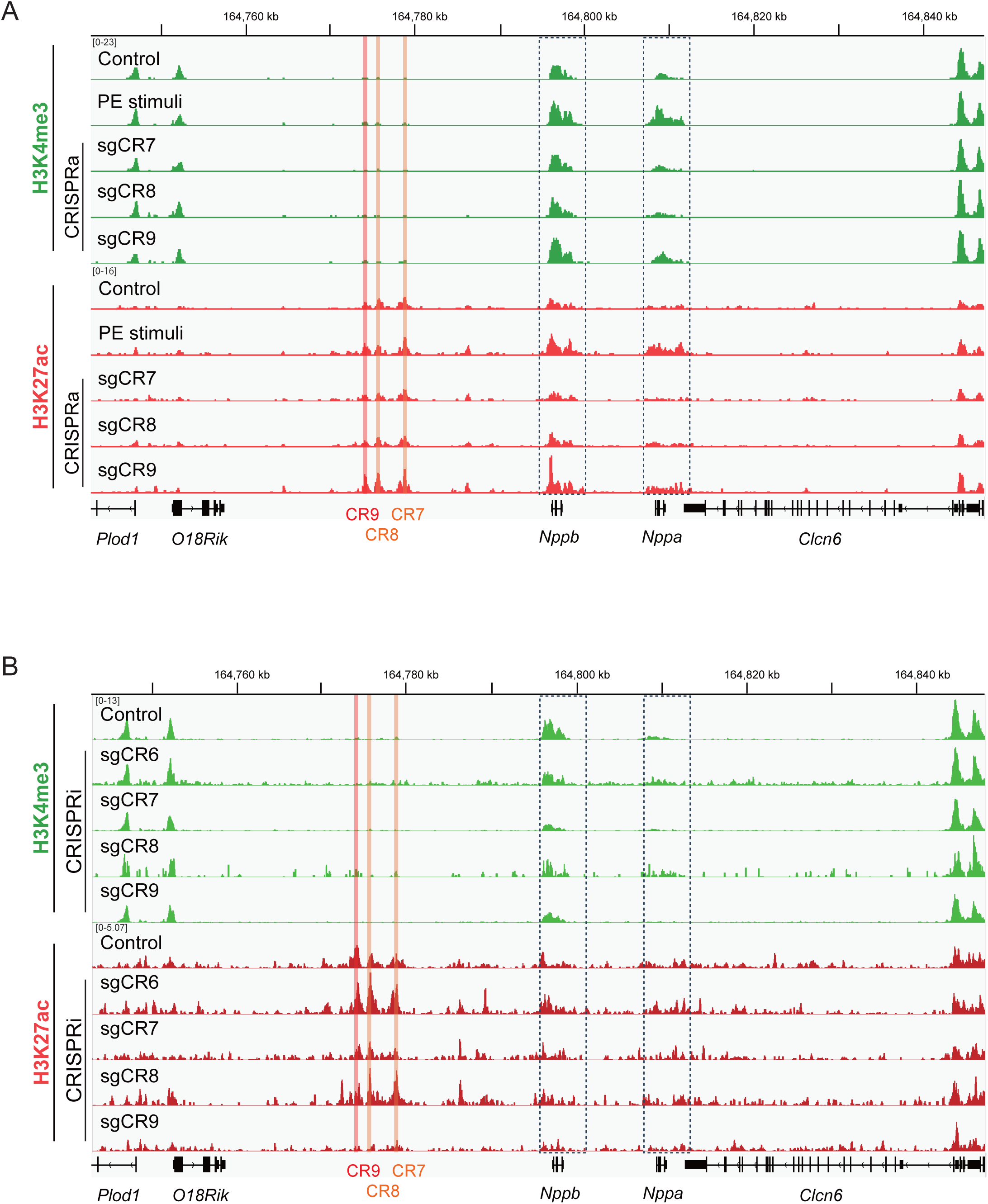
CRISPR-based CR9 modulation remodels chromatin accessibility and histone modifications across the RE1 enhancer cluster and the *Nppa–Nppb* promoters. **(A)** Density maps show H3K4me3 (green) and H3K27ac (red) ChIP–seq profiles in neonatal rat cardiomyocytes following CRISPRa targeting of CR7, CR8, or CR9, or stimulation with phenylephrine (PE), which serves as a positive control for *Nppa* and *Nppb* induction. Data are shown across chr5:164,743,000– 164,848,000 (rn6). CR9 region is marked in red and CR7/CR8 in orange. Dotted boxes indicate the *Nppa* and *Nppb* gene loci. **(B)** H3K4me3 and H3K27ac ChIP–seq profiles as in (A), but following CRISPRi targeting of CR6–CR9. All sequencing experiments were performed in two biologically independent replicates and confirmed in at least two separate experiments.

Conversely, CRISPRi of CR9 resulted in marked reductions in H3K27ac signals across the CR7–CR9 region and decreased levels of both H3K4me3 and H3K27ac at the *Nppa/Nppb* promoters (Figure 2B). In contrast, CRIPSRa or CRISPRi targeting CR7 or CR8 did not substantially alter the histone modifications in the enhancer or promoter regions. These findings support a causal role for CR9 in maintaining chromatin accessibility and promoter activity, likely through enhancer–promoter interactions that modulate histone modifications at target gene promoters. Notably, chromatin states in each organ indicated that CR9 was inactive in non-cardiac cell types, underscoring its tissue-specific regulatory role (Supplementary Figure 3).

### CR9 cooperates with neighboring enhancers and spatially associates with the *Nppb* transcription site

To explore the functional interactions between the three constituent enhancers (CR7, CR8, and CR9), we performed combinatorial activation and inhibition experiments while keeping the viral load constant. Compared to the activation of CR9 alone, co-activation of CR9 with either CR7 or CR8 resulted in synergistic upregulation of *Nppa* and *Nppb* mRNA expression (Figure 3A). Conversely, while individual CR9 inhibition significantly reduced *Nppa* and *Nppb* expression, the simultaneous inhibition of CR9, CR7, and CR8 further decreased *Nppa* and *Nppb* expression (Figure 3B).

**Figure 3.**
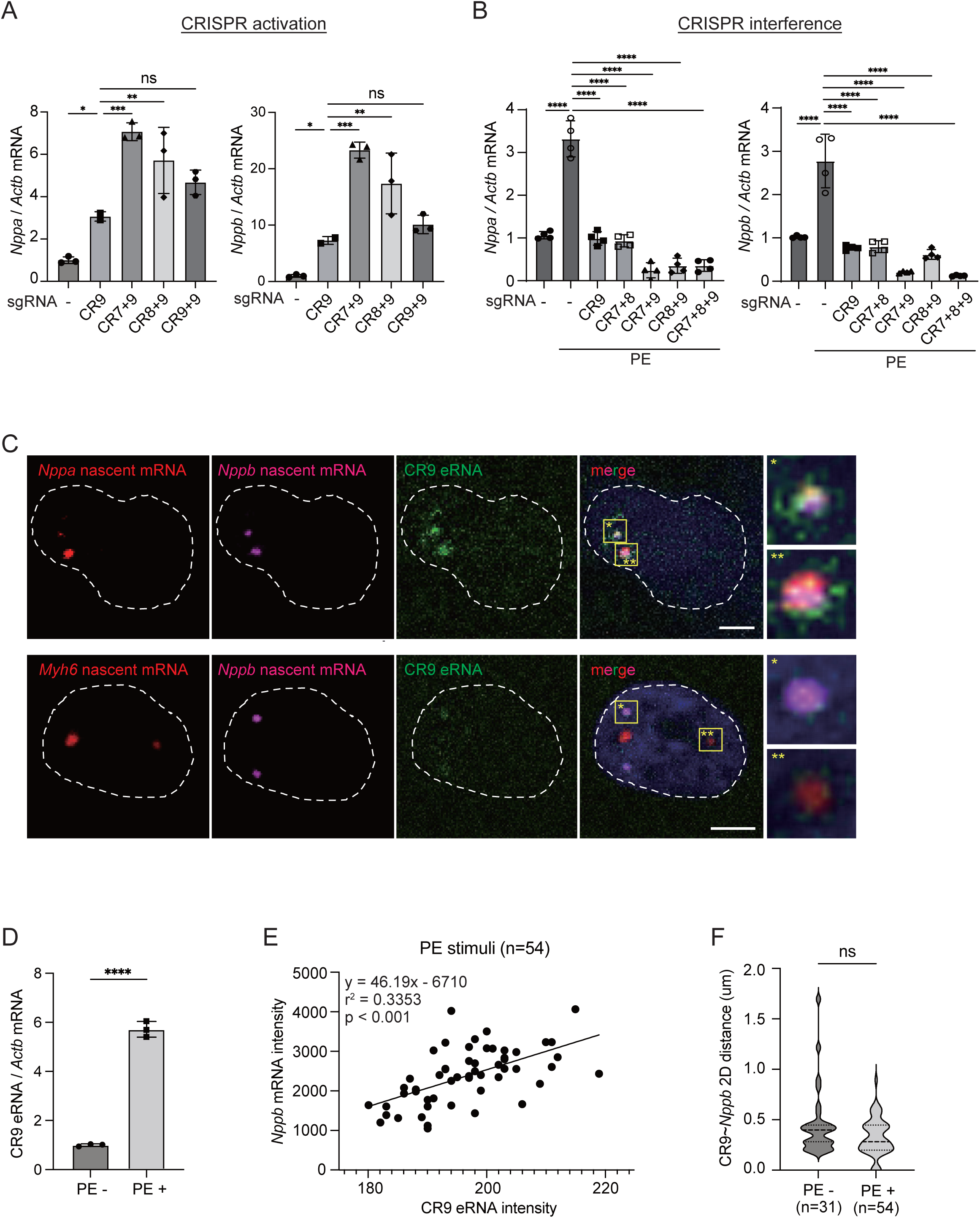
CR9 functions as a regulatory hub that is both required and sufficient to induce transcription and spatial coordination of *Nppa* and *Nppb* expression. **(A)** qPCR analysis of *Nppa* and *Nppb* expression in neonatal rat cardiomyocytes following CRISPRa targeting of CR7, CR8, CR9, or combinations thereof. Expression was normalized to *Actb* and shown relative to sgCtrl. **(B)** As in (A), but following CRISPRi targeting of CR7, CR8, or CR9, with or without PE stimulation. **(C)** Representative RNA-FISH images showing nuclear signals for CR9 eRNA (green), *Nppb* nascent RNA (red), and either *Nppa* (yellow, top) or *Myh6* (yellow, bottom) nascent transcripts. Scale bar, 3 μm. **(D)** CR9 eRNA expression levels in neonatal rat cardiomyocytes under starvation or PE stimulation. **(E)** Correlation between CR9 eRNA and *Nppb* nascent transcript intensities in PE-treated cardiomyocytes, based on RNA-FISH signals shown in (C). **(F)** Violin plot showing the 2D distances between CR9 eRNA and *Nppb* transcription sites under starvation or PE stimulation, based on RNA-FISH signals shown in (C). All individual data points are shown. Data represent mean ± s.d.; *n* = 4 biologically independent experiments for (A, B), and 3 for (D); *n* = 31 (without PE) and 54 (with PE) nuclei for RNA-FISH analysis (C, E, and F). Statistical significance was determined using one-way ANOVA with Tukey’s post hoc test (A, B), unpaired two-tailed Student’s t-test (D, F), and linear regression analysis (E). **** *P* < 0.0001 (E); *** *P* < 0.001 (A, B); ** *P* < 0.01 (A, B); * *P* < 0.05 (A, B, F); *P* = 0.0324 (F); *r*² = 0.3353, *P* < 0.001 (E); ns, not significant.

To assess the spatial relationship between enhancer and promoter activities, we visualized the CR9-derived enhancer RNA (eRNA) and nascent transcripts of *Nppa* and *Nppb* using RNAscope^22^ in cultured rat cardiomyocytes. These three signals were frequently observed in proximity within the nucleus (Figure 3C). In contrast, CR9 eRNA showed no detectable co-localization with nascent *Myh6* transcripts, indicating that its spatial association was specific to the *Nppa–Nppb* locus (Figure 3C). Phenylephrine (PE) stimulation, known to induce the expression of *Nppa* and *Nppb,* increased the fluorescence intensity of CR9 eRNA, which positively correlated with *Nppb* transcription levels (Figure 3D and 3E), reflecting enhanced enhancer activity under stress conditions.

Notably, the two-dimensional spatial distances between the respective foci remained unchanged (Figure 3F), suggesting that transcriptional activation may occur without further reduction in physical proximity. This observation is consistent with recent reports that enhancer–promoter distances may remain stable or even increase during active transcription^23–25^, supporting a transcriptional hub model based on multivalent phase-separated condensates^26–29^.

### CR9 is required for *Nppa–Nppb* expression in human iPSC-derived cardiomyocytes

To examine whether *Nppa–Nppb* expression is also dependent on CR9 in human cardiomyocytes, we deleted the CR9 element in human iPSC-derived cardiomyocytes using active Cas9 protein and two sgRNA targeting sequences within the CR9 region. We identified a single iPSC clone in which both the CR9 enhancer alleles were successfully deleted, as confirmed by PCR genotyping and Sanger sequencing (Supplementary Figure 4A and 4B). This homozygous ΔCR9 clone was subsequently used for downstream expression analyses.

Deletion of the CR9 region resulted in a marked reduction in *Nppa* and *Nppb* mRNA levels (Supplementary Figure 4C). These findings confirm that CR9 is essential for proper *Nppa–Nppb* expression and functions as a critical enhancer element in human cardiomyocytes.

### CR9 deletion disrupts *Nppa–Nppb* transcription and chromatin activation in vivo

To evaluate the in vivo requirement of CR9 for *Nppa*/*Nppb* expression, we generated heterozygous and homozygous CR9 knockout mice (CR9⁺/⁻ and CR9⁻/⁻) using homologous recombination to delete the CR9 element (Supplementary Figures 5A to 5C). These mice were viable and fertile, with no overt phenotypes other than mild hypertension during the first few postnatal months. Transcriptomic analysis of the LV tissue from 8-week-old CR9⁻/⁻ and wild-type (WT) mice revealed marked downregulation of *Nppa* and *Nppb* in CR9⁻/⁻, with intermediate reduction in CR9⁺/⁻ mice, as confirmed by qRT-PCR (Figure 4A). In contrast, at embryonic day 16 (E16) and postnatal day 1(P1), *Nppa* and *Nppb* levels in CR9⁻/⁻ mice were comparable to WT (Figure 4B), suggesting a developmental switch in enhancer usage, with CR9 becoming functionally important after birth.

**Figure 4.**
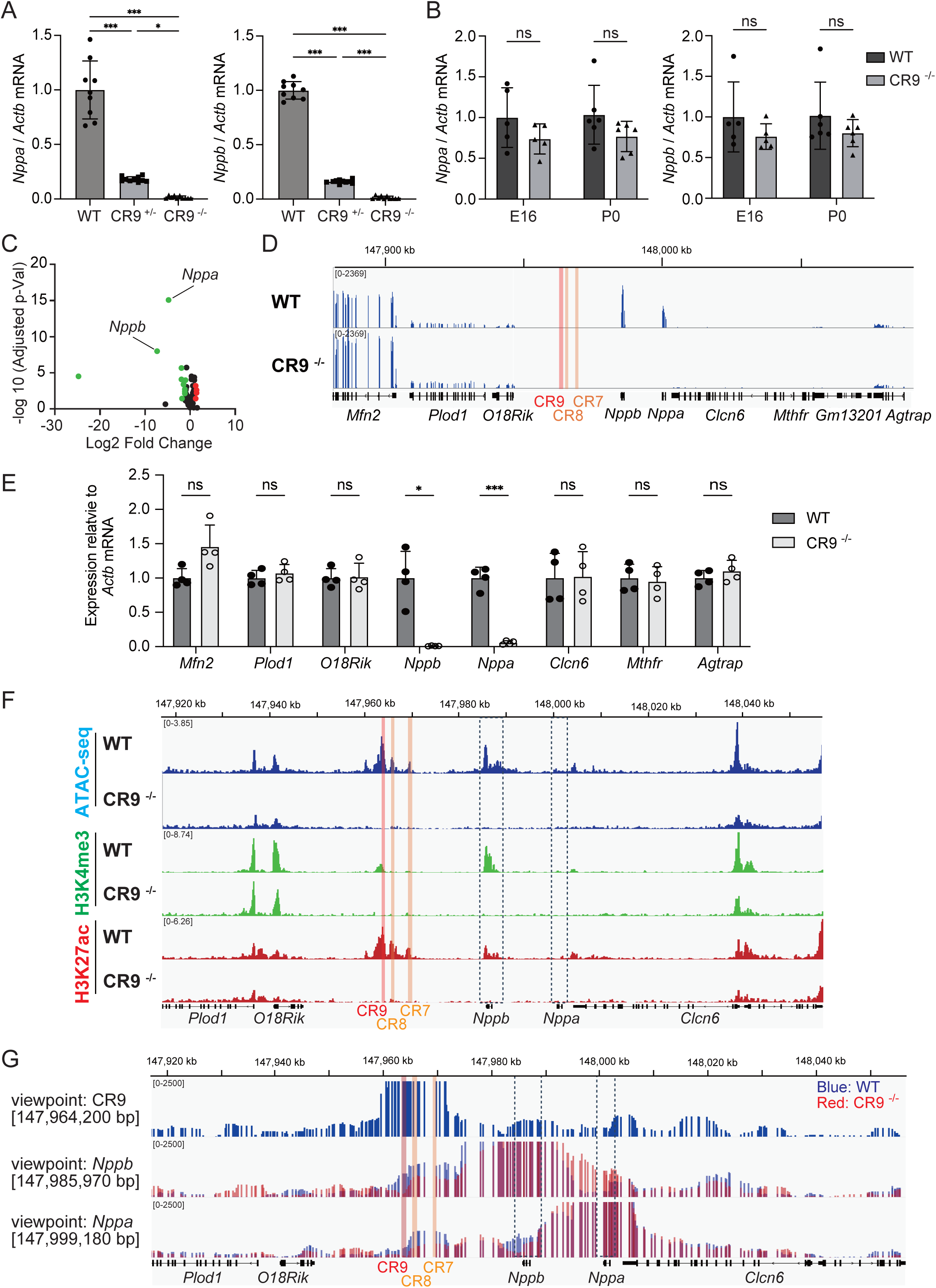
CR9 deletion reduces *Nppa* and *Nppb* expression and chromatin activation in vivo. **(A, B)** qPCR of *Nppa* and *Nppb* mRNA in left ventricles (LVs) from wild-type (WT) and CR9⁻/⁻ male mice at 8 weeks of age (A), and at embryonic day 16 (E16) and postnatal day 1 (P1) (B). CR9⁺/⁻ mice were also included in (A). Expression was normalized to *Actb* and shown relative to WT at each time point. **(C)** Volcano plot showing differentially expressed genes in LVs of CR9^-/-^ mice compared to WT, based on RNA-seq data. Genes with false discovery rate (FDR) < 0.05 and log₂ fold change > 1 are shown in color: red for upregulated and green for downregulated genes. *Nppa* and *Nppb* were significantly downregulated. **(D)** Representative RNA-seq coverage tracks in the *Nppa–Nppb* locus (chr4: 147,881,298–148,151,825; mm10) and flanking genes in LVs of WT and CR9*^-/-^* mice. CR9 region is marked in red; CR7 and CR8 in orange. Dotted boxes indicate the *Nppa* and *Nppb* loci. **(E)** qPCR analysis of *Nppa*, *Nppb*, and the neighboring genes in LVs from WT and CR9^-/-^ male mice. Expression was shown relative to WT. **(F)** Density maps of chromatin accessibility (ATAC-seq, blue), H3K4me3 ChIP-seq (green), and H3K27ac ChIP-seq (red) signals across the *Nppa–Nppb* locus (chr4:147,917,000–148,057,000; mm10) in LVs from WT and CR9^⁻/⁻^ mice. Enhancer regions CR9 (red), CR7 and CR8 (orange), and gene loci (dotted boxes) are indicated. **(G)** Three circular Chromosome Conformation Capture (4C) contact maps from LVs of WT and CR9^⁻/⁻^ mice are shown using the following viewpoints: CR9 (top panel), the *Nppb* promoter (middle panel), and the *Nppa* promoter (bottom panel). Contact frequencies are shown for WT (blue) and CR9^⁻/⁻^ (red) hearts. Data represent mean ± s.d.; *n* = 9 to 10 for (A), 5 to 6 for (B), and 4 for (D). Statistical significance was determined using one-way ANOVA (A), two-way ANOVA (D), and FDR-corrected differential expression analysis (B). *** P < 0.001 (D); ** P < 0.01 (A); * P < 0.05 (A); ns, not significant.

RNA-seq identified 15 significantly differentially expressed genes (fold change > 2, FDR < 0.05) in CR9⁻/⁻ LV tissue (Figure 4C and Supplementary Table 3), while neighboring genes such as *Miip*, *Mfn2*, *Plod1*, and *KIAA2013* were unaffected (Figure 4D), as confirmed by qPCR (Figure 4E). These results indicate that CR9 specifically regulates the *Nppa–Nppb* locus in vivo with minimal impact on nearby genes.

In the WT LV tissue, ATAC-seq revealed three major chromatin accessibility peaks, corresponding to CR7, CR8, and CR9, in addition to the peaks at the promoters and gene bodies of *Nppa* and *Nppb* (Figure 4F). These regions also exhibited strong enrichment of H3K27ac and H3K4me3 histone markers (Figure 4F). In CR9⁻/⁻ mice, both chromatin accessibility and active histone marks were markedly reduced not only at the CR9 site, but also at CR7/8 and the promoters and gene bodies of *Nppa* and *Nppb*. These findings suggest that CR9 is essential for maintaining a transcriptionally active chromatin landscape at the *Nppa–Nppb* locus in vivo.

To further evaluate whether CR9 directly regulates its target promoters via chromatin looping in vivo, we performed 4C-seq using nuclei purified from LV tissues of WT and CR9⁻/⁻ mice. Viewpoint-specific primers were designed to quantify DNA fragments interacting with the promoters of *Nppa* and *Nppb*. In WT hearts, the CR9 element showed strong physical interactions with both *Nppa* and *Nppb* promoters simultaneously (Figure 4G). In contrast, such interactions were absent in CR9⁻/⁻ mice, and the interaction frequencies between the promoters and other enhancer elements (CR7 and CR8) were also markedly reduced (Figure 4G). These results suggest that CR9 plays a pivotal role in mediating enhancer–promoter engagement at the *Nppa–Nppb* locus in vivo, serving as a central regulatory hub within the SE structure.

### CR9 knock-in at the native locus restores *Nppa–Nppb* expression and chromatin activity

To confirm the functional importance of CR9 in vivo, we generated additional mouse lines by combining CRISPR-Cas9 with single-stranded oligodeoxynucleotides (ssODNs)^30^ encoding the CR9 element. One line carried a knock-in of the CR9 sequence at its native locus in the forward orientation (CR9^Fw/Fw^) and the other in the reverse orientation (CR9^Rv/Rv^) (Figure 5A, Supplementary Figure 6A). CR9^Fw/Fw^ mice showed complete restoration of *Nppa* and *Nppb* expression (Figure 5B), as well as the recovery of chromatin accessibility and active histone modifications at the locus (Figure 5C). In contrast, mice carrying a control ssODN sequence derived from the CR3 region did not exhibit a rescue of gene expression or chromatin features (Figures 5B and 5C), supporting the element-specific enhancer activity of CR9. Interestingly, CR9^Rv/Rv^ mice also displayed restored gene expression and chromatin activity, consistent with the orientation-independent nature of enhancer function^31^.

**Figure 5.**
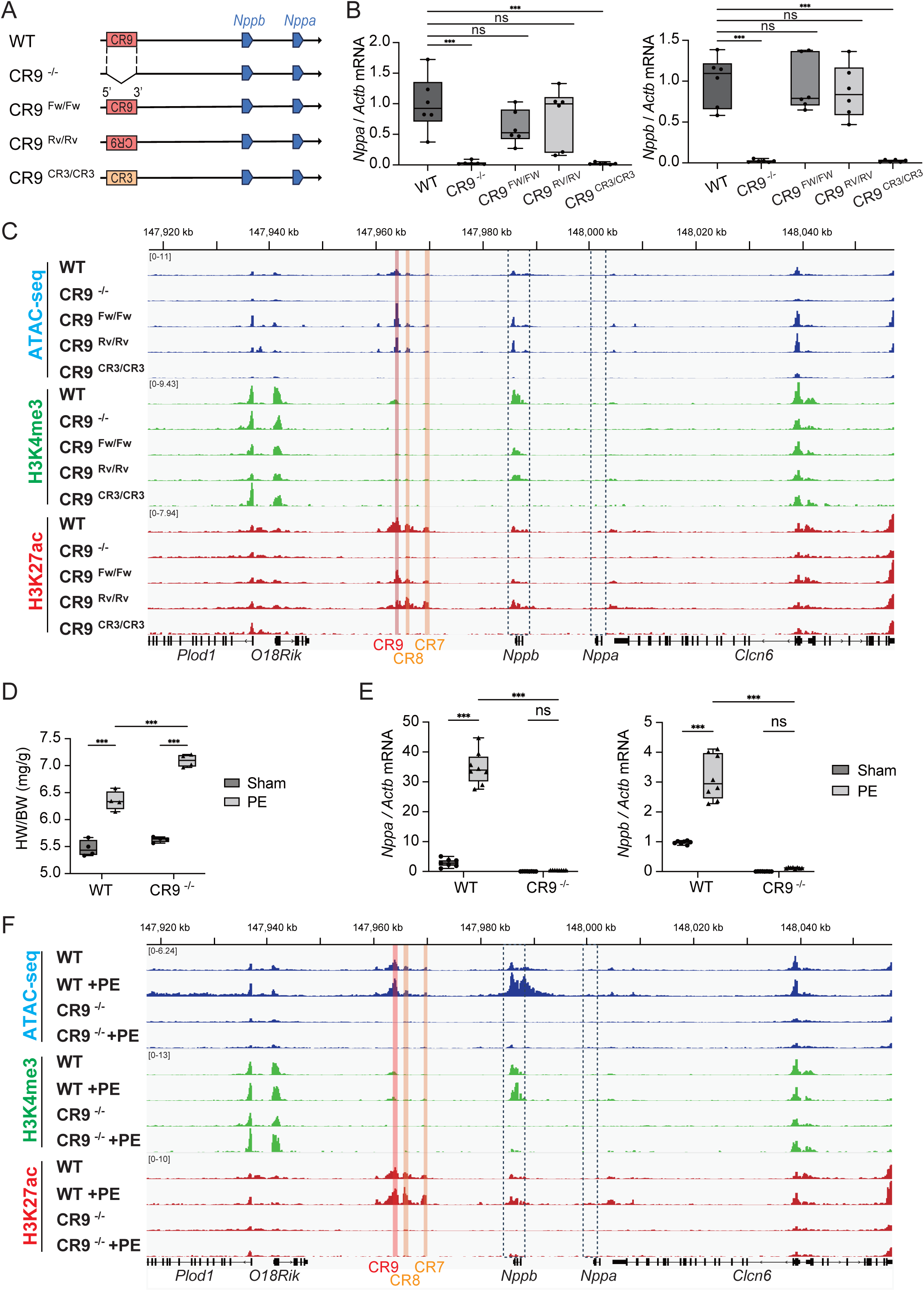
CR9 knock-in restores transcription and chromatin state at *Nppa-Nppb* locus. **(A)** Schematic representations of the CR9 knockout and rescue strategy in mice. The CR9 forward (CR9^Fw/Fw^), CR9 reverse (CR9^Rv/Rv^) or a control CR3 sequence (CR9^CR3/CR3^) was reintroduced into the endogenous CR9 locus in CR9^-/-^ mice. **(B)** qPCR analysis of *Nppa* and *Nppb* mRNA in LV myocardium from WT, CR9^-/-^, CR9^Fw/Fw^, CR9^Rv/Rv,^ and CR9^CR3/CR3^ male mice. Expression is shown relative to WT. **(C)** Density maps of ATAC-seq (blue), H3K4me3 ChIP-seq (green), and H3K27ac ChIP-seq (red) signals across the *Nppa-Nppb* loci (chr4:147,917,000–148,057,000; mm10) in LVs from WT, CR9^⁻/⁻^, CR9^Fw/Fw^, CR9^Rv/Rv^, and CR9^CR3/CR3^ mice. Enhancer regions CR9 (red), CR7 and CR8 (orange), and gene loci (dotted boxes) are indicated. **(D)** Heart weight to body weight (HW/BW) ratio in 9-week-old male mice with or without PE treatment for 7 days. **(E)** *Nppa* and *Nppb* mRNA levels in LV myocardium of WT and CR9^-/-^ male mice treated with or without PE for 7 days. Expression was normalized to the untreated group (set as 1). **(F)** Density maps of ATAC-seq, H3K4me3 ChIP-seq, and H3K27ac ChIP-seq signals across the *Nppa–Nppb* locus in WT and CR9^⁻/⁻^ mice treated with or without PE, as in (C). Data represent mean ± s.d.; *n* = 6 for (B), 4 for (D), and 7 for (E). Statistical significance was determined using one-way ANOVA (B), two-way ANOVA (D, E), followed by Fisher’s least significant difference (LSD) test for (E). *** *P* < 0.001 (B, D, E); ** *P* = 0.01 (E); ns, not significant.

To investigate the role of CR9 under pathological conditions, we subjected 8-week-old CR9⁻/⁻ and WT mice to 7-day PE treatment, which is known to induce pathological cardiac hypertrophy. Both groups developed significant LV hypertrophy after PE administration (Figure 5D and Supplementary Figure 7A and 7B). Although PE stimulation significantly upregulated *Nppa* and *Nppb* expression in WT hearts, this response was not statistically significant in CR9⁻/⁻ mice. While fold-change values appeared even greater in CR9⁻/⁻ hearts, this was due to markedly reduced baseline expression, leading to inflated ratios. In absolute terms, PE-treated CR9⁻/⁻ hearts failed to reach the expression levels observed in WT counterparts. Consistent with impaired induction of *Nppa* and *Nppb*, the degree of PE-induced hypertrophy was more pronounced in CR9⁻/⁻ mice (Figure 5D), supporting the antihypertrophic role of natriuretic peptides in this context. These results indicate that CR9 deficiency alters transcriptional responsiveness to hypertrophic stimuli (Figure 5E).

Next, we assessed chromatin accessibility and histone modification states at the *Nppa–Nppb* locus using ATAC-seq and ChIP-seq, respectively. In WT hearts, PE stimulation increased chromatin accessibility at the CR9 region and around the *Nppb* gene (Figure 5F), accompanied by enrichment of the activating histone marks H3K27ac and H3K4me3. In contrast, CR9⁻/⁻ mice failed to exhibit these epigenetic changes in response to PE stimulation. Taken together, these results demonstrate that CR9 functions as a key enhancer regulating *Nppa* and *Nppb* expression under both basal and stress-induced conditions. Notably, these regulatory and epigenetic changes were confined to *Nppa– Nppb* TAD, with no detectable alterations observed outside the domain, consistent with a topologically restricted enhancer function.

### CR9 chromatin accessibility is dynamically regulated by mechanical unloading in human heart failure

Finally, we investigated chromatin state changes within the *NPPA–NPPB* locus in LV tissue from patients with severe heart failure who underwent LVAD implantation (Figure 6A and Supplementary Table 4). We analyzed two paired samples from patients with dilated cardiomyopathy who exhibited markedly elevated plasma BNP levels and impaired hemodynamics prior to surgery, both of which improved following LVAD implantation (Figure 6A). Echocardiographic images before and after LVAD implantation demonstrated a reduction in the LV chamber size, visually supporting effective hemodynamic unloading (Figure 6B and 6C). qPCR confirmed marked reductions in *NPPA* and *NPPB* mRNA expression in LV tissues after LVAD implantation in both patients (Figure 6D). Intriguingly, ATAC-seq revealed that chromatin accessibility at the CR9 region and across the *NPPA–NPPB* locus was markedly decreased after mechanical unloading by the LVAD (Figure 6E). These findings suggest that CR9 activity is dynamically regulated by cardiac load in vivo and that the enhancer–promoter axis at this locus is responsive to mechanical stress, even in the failing human heart.

**Figure 6.**
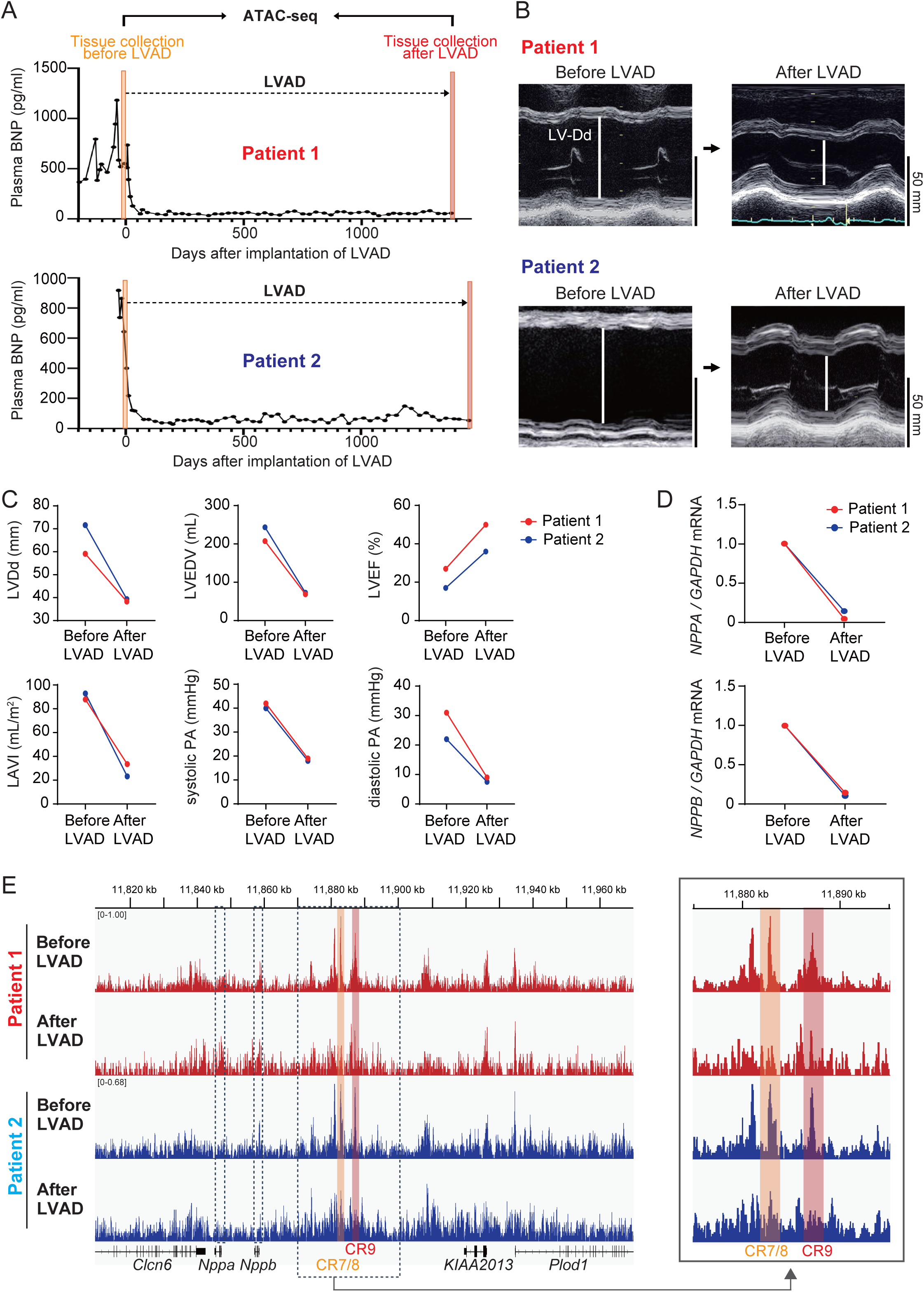
Chromatin accessibility surrounding CR9 is dynamically altered by LVAD-mediated unloading in human heart failure. **(A)** Time-course of plasma B-type natriuretic peptide (BNP) levels in two patients with dilated cardiomyopathy (DCM) before and after implantation of a left ventricular assist device (LVAD). Day 0 indicates the timing of LVAD surgery. Longitudinal BNP levels in Patient 1 (top) and Patient 2 (bottom) are shown. Orange bars indicate myocardial samples collected before LVAD (high BNP); red bars indicate samples collected after LVAD (low BNP). **(B)** Hemodynamic and echocardiographic parameters in Patients 1 and 2 before and after LVAD implantation, including left ventricular end-diastolic diameter (LVDd), end-diastolic volume (LVEDV), ejection fraction (LVEF), left atrial volume index (LAVI), and pulmonary artery pressures (systolic and diastolic). Red and blue lines represent Patients 1 and 2, respectively. **(C)** qPCR analysis of *NPPA* and *NPPB* mRNA levels in LV myocardium from Patients 1 (red) and 2 (blue) before and after LVAD implantation. Expression was normalized to *GAPDH*. **(D)** ATAC-seq signal intensity across the *NPPA–NPPB* locus (chr1:11,810,000–11,970,000; hg38) in LV myocardium before and after LVAD implantation. Enhancer regions CR7 (red), CR8, and CR9 (orange) are indicated. Red and blue traces correspond to Patients 1 and 2, respectively.

## Discussion

In this study, we identify CR9 element as a stress-inducible hub enhancer within the *Nppa*/*Nppb* super-enhancer, orchestrating the transcription of *Nppa* and *Nppb*, the canonical biomarkers and cardioprotective hormones of heart failure. We show that CR9 is both necessary and sufficient for the robust induction of natriuretic peptides under pathological stress. Importantly, by analyzing paired myocardial samples from the same patients before and after LVAD unloading, we provide the first direct evidence that chromatin accessibility at the CR9 element is dynamically reversible in human heart failure, underscoring enhancer plasticity as a fundamental property of the diseased myocardium. These findings establish the molecular basis of natriuretic peptide regulation, highlight a hierarchical organization within the *Nppa*/*Nppb* super-enhancer, and reveal enhancer plasticity as a potential therapeutic axis.

The *Nppa/Nppb* locus provides a powerful model to dissect enhancer hierarchy and multi-gene regulation within a SE context. It is embedded within a well-insulated TAD bounded by convergent CTCF sites and contains two adjacent stress-inducible genes, *Nppa* and *Nppb*, that are coordinately activated under pathological conditions together with an SE composed of multiple constituent elements. Our functional dissection revealed that CR9 is the dominant hub enhancer within this locus and displays several defining features. First, CR9 establishes a functional hierarchy: although multiple conserved elements reside within RE1, CR9 exerts dominant regulatory control, while CR7 and CR8 show only modest activity. In line with observations from other SEs, dominant hub elements can coordinate both transcription and chromatin structure within TADs^9,15,16,32^, providing a spatial framework for specificity and robustness in gene regulation. Second, CR9 is necessary for natriuretic peptide induction, as CRISPRi or deletion markedly reduced *Nppa*/*Nppb* expression. Third, CR9 is sufficient to drive transcription, since CRISPRa stimulation or reinsertion of CR9 alone restored target gene activation. Fourth, CR9 functions cooperatively with neighboring enhancers, because co-activation with CR7 or CR8 yielded synergistic transcription despite their weak independent activity. These results demonstrate that enhancer elements with similar chromatin marks can differ in their transcriptional impact^33,34^, emphasizing the importance of functional dissection. Fifth, CR9 exhibits orientation-independent activity, as reinsertion in reverse orientation reinstated gene expression and chromatin accessibility. A similar orientation-independent behavior has been reported for the major enhancer R2 within the α-globin SE^35^. Finally, the regulatory influence of CR9 remained spatially confined within the boundaries of the TAD, indicating that CR9 acts as a locus-specific enhancer rather than broadly perturbing the surrounding genome. Together, these findings establish CR9 as a dominant hub enhancer that integrates hierarchical, cooperative, and structural features to coordinate natriuretic peptide expression under cardiac stress.

Moreover, we uncovered that the *Nppa/Nppb* enhancer landscape undergoes a developmental switch: CR6 and CR7 are active in the embryonic heart^36^ but are silenced postnatally (Supplementary Figure 8), whereas CR9 becomes increasingly accessible and acetylated after birth and responds to mechanical stress. This suggests that CR9 acts as a latent enhancer^37^, activated by the hemodynamic load. ATAC-seq and ChIP-seq revealed minimal chromatin accessibility and H3K27ac at CR9 during development, which increased postnatally in parallel with cardiac pressure, which is consistent with its previously reported load-inducible function^2,3^. While our previous studies showed only the correlative activation of CR9 and *Nppa/Nppb*^2,3^, our current data established their causal and hierarchical roles in chromatin remodeling and transcriptional control. Together, these findings highlights a regulatory switch from developmental enhancer to a stress-inducible enhancer^37–39^, positioning CR9 as a key regulatory element of natriuretic peptide expression in the adult heart (Figure 7).

**Figure 7.**
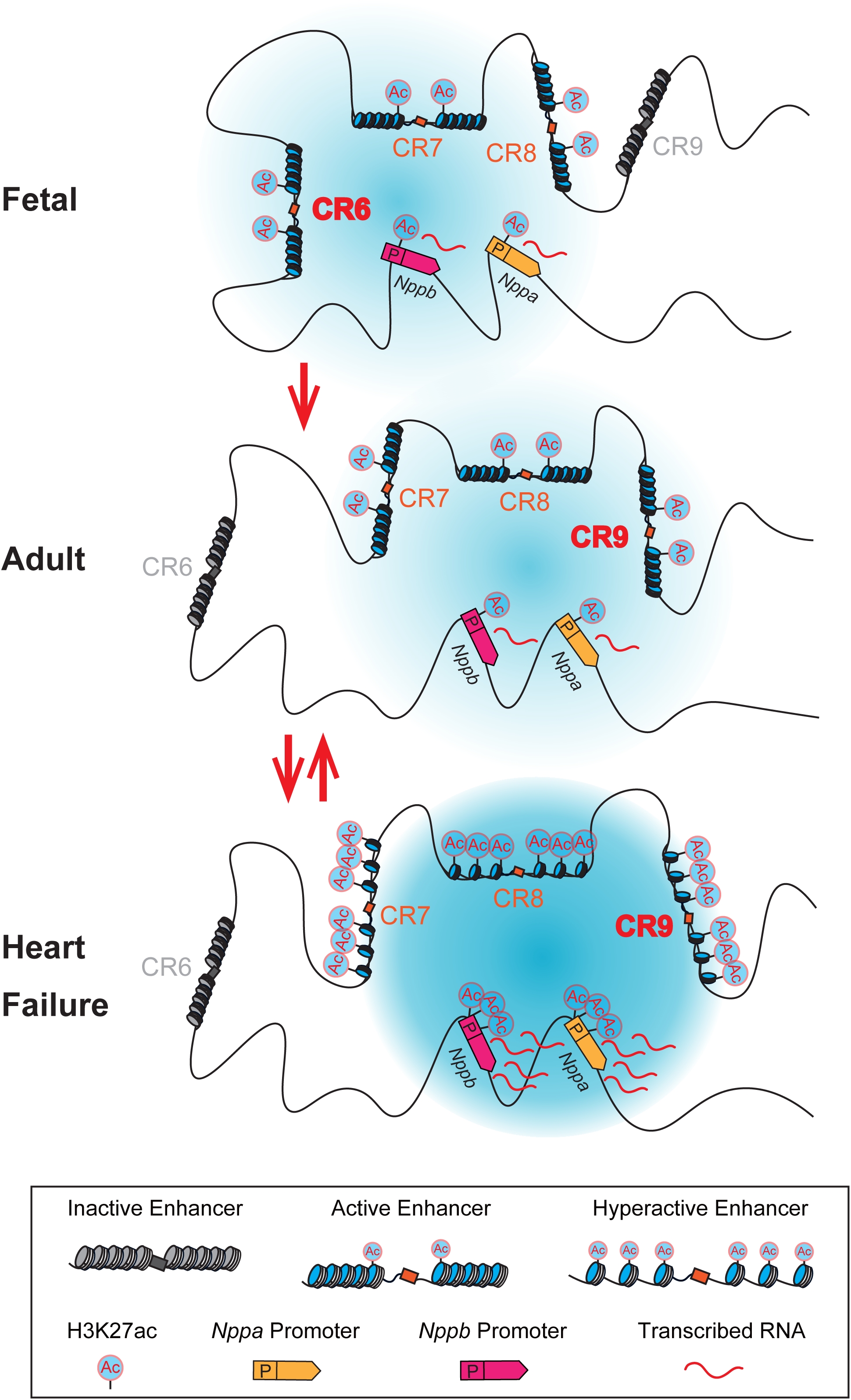
Conceptual model of dynamic enhancer usage at the *Nppa/Nppb* locus during heart development and disease. Schematic representation of enhancer activity surrounding the *Nppa* and *Nppb l*oci in fetal, adult, and failing hearts. During fetal development, CR6 is proposed to act as the primary enhancer regulating natriuretic peptide gene expression, based on previous studies^36^. In the adult heart, the enhancer landscape is reorganized, and CR9 emerges as a dominant hub enhancer coordinating the transcription of *Nppa* and *Nppb*. In heart failure, CR9 is further activated in response to hemodynamic stress and contributes to the pathological upregulation of *Nppa* and *Nppb*.

Active enhancers are known to recruit transcription factors and RNA polymerase II and generate short, unstable enhancer RNAs (eRNAs) that serve as hallmarks of enhancer activity^40,41^. In the current study, CR9-derived eRNA increased in response to hypertrophic stress and correlated with nascent *Nppa*/*Nppb* transcripts, indicating that eRNA production at CR9 is tightly coupled to transcriptional output at its target promoters. These findings support models in which enhancer transcription precedes and facilitates promoter activation^41^, extending this paradigm to the cardiac SE context. Beyond classical looping models^7^, recent work has proposed a transcription hub model^9,26,27,29,42^ in which transcription factors and RNA polymerase II form multivalent condensates that can simultaneously engage multiple promoters^26,43,44^, even at relatively large spatial distances. In line with this concept, we found that CR9 physically interacted with both *Nppa* and *Nppb* promoters in wild-type hearts, whereas these contacts were lost in CR9 knockout. Moreover, RNAscope revealed that CR9 eRNA co-localized with both *Nppa* and *Nppb* nascent transcripts, with proximity maintained during active transcription^23–25^, consistent with the transcription hub model^4,27,39^ in which multivalent condensates enable enhancer–promoter communication without stable looping. These observations suggest that CR9 functions within a transcription hub, organizing cooperative rather than competitive^4^ enhancer–promoter interactions to coordinate natriuretic peptide gene activation under stress. Nonetheless, the coordinated activation of *Nppa* and *Nppb* upon CR9 perturbation supports a model in which CR9 orchestrates multigene transcription, favoring cooperative enhancer– promoter communication over competition.

The most important finding of this study is that we provide the first direct evidence that enhancer states are dynamically reversible in the human failing heart. In paired ventricular samples obtained before and after LVAD unloading, we observed that chromatin accessibility at the CR9 element was restored with relief of hemodynamic stress. This demonstrates that enhancer activation in failing myocardium is not permanently fixed but can be reset by therapeutic unloading. Such reversibility offers a molecular explanation for the well-known decline in natriuretic peptide levels with effective therapy and highlights enhancer plasticity as a disease-relevant property of the failing heart. Because *NPPA* and *NPPB* encode both indispensable biomarkers and cardioprotective hormones, the identification of CR9 as their regulatory hub establishes a direct mechanistic link between hemodynamic stress and clinical practice. By fine-mapping the broad RE1 region to a 650-bp segment (CR9) with definitive enhancer function, our study delineates the minimal regulatory unit required for stress-inducible natriuretic peptide expression. Ongoing work is further narrowing this region to identify the core sequence sufficient to act as an autonomous hub enhancer. Acting as a sensor of hemodynamic stress, CR9 activates downstream gene expression programs, exemplified by the induction of natriuretic peptides. Such enhancer plasticity offers a conceptual framework for developing gene therapy strategies in which CR9 controls the conditional expression of therapeutic transgenes in failing hearts. By sensing mechanical load and inducing the expression of cardioprotective natriuretic peptides, CR9 may contribute to adaptive responses during heart failure progression. Its reversibility and conservation across humans, mouse, and iPSC-derived cells highlight CR9’s potential as a therapeutic target for heart failure and related stress-responsive diseases. In the future, synthetic constructs incorporating the core sequence of the CR9 enhancer could enable stress-responsive cardiac gene therapies, driving protective genes expression in proportion to myocardial stress.

These principles may be extended across other regulatory systems, providing a conceptual basis for enhancer-targeted therapies in cardiovascular and other stress-associated diseases. This enhancer-centric framework not only elucidates the molecular basis of natriuretic peptide regulation but also provides a blueprint for translating enhancer biology into therapeutic strategies for heart failure. These insights bridge molecular mechanisms with bedside biomarkers, offering a direct conceptual link between enhancer biology and heart failure management.

This study has several limitations. First, our human data are limited to two paired pre- and post-LVAD heart failure cases and should be considered exploratory. Nevertheless, these rare paired samples provide a unique opportunity to directly track enhancer dynamics in response to hemodynamic unloading in human tissues, an analysis that is rarely achievable in clinical practice. Future studies in larger patient cohorts will be required to validate and extend these findings. Second, although CR9 emerged as a dominant hub enhancer, our current chromatin conformation assays cannot resolve whether CR9 engages the *Nppa* and *Nppb* promoters simultaneously or sequentially. Techniques such as 4C-seq and RNAscope demonstrate physical proximity and nuclear co-localization, but they lack the resolution to establish whether CR9 engages both promoters concurrently within the same nucleus. Future studies employing higher-resolution techniques, such as live-cell imaging^43,45^ or dual-probe RNA labeling^29,30^, will be required to determine the precise dynamics of CR9–promoter communication. Finally, although our findings provide a mechanistic example of stress-inducible enhancer function in vivo, it remains unclear whether the hierarchical and cooperative enhancer models observed at the *Nppa/Nppb* locus are broadly applicable across cardiac loci. For example, while *Nppa* and *Nppb* are co-induced under stress, the *Myh6/Myh7* cluster shows reciprocal expression controlled by a shared but mutually exclusive enhancer^46^, suggesting that hub enhancer logic may be locus-specific. Further studies in other stress-responsive systems, and potentially in other tissues such as skeletal muscle or immune cells, will be required to clarify the generalizability of this regulatory architecture.

## Data Availability

The sequencing data generated in this study will be deposited in the GEO database prior to publication. All the other data are available from the corresponding author upon request.

## Abbreviations

ANP: atrial natriuretic peptide
ATAC-seq: assay for transposase-accessible chromatin using sequencing
BNP: B-type natriuretic peptide
ChIP-seq: chromatin immunoprecipitation sequencing
CRISPRa: CRISPR activation
CRISPRi: CRISPR interference
CTCF: CCCTC-binding factor
DAPI: 4′,6-diamidino-2-phenylindole
dCas9: dead Cas9 (catalytically inactive Cas9)
eRNA: enhancer RNA
ES cell: embryonic stem cell
FBS: fetal bovine serum
FDR: false discovery rate
gRNA: guide RNA
H3K27ac: histone H3 lysine 27 acetylation
H3K4me3: histone H3 lysine 4 trimethylation
iPSC: induced pluripotent stem cell
LV: left ventricle / left ventricular
LVAD: left ventricular assist device
MC1-DTA: diphtheria toxin A under MC1 promoter
NPPA: natriuretic peptide A (atrial natriuretic peptide gene)
NPPB: natriuretic peptide B (B-type natriuretic peptide gene)
PBS: phosphate-buffered saline
PE: phenylephrine
PGK-Neo: phosphoglycerate kinase–neomycin resistance cassette
qPCR: quantitative polymerase chain reaction
RNA-seq: RNA sequencing
RT-PCR: reverse transcription polymerase chain reaction
SE: super-enhancer
sgRNA: single guide RNA
ssODN: single-stranded oligodeoxynucleotide
TAD: topologically associating domain

## Acknowledgments

This work was supported by grants from the Japan Society for the Promotion of Science KAKENHI [20K17151, 22K08204, 23K24327, 25K11386], by grants from the Japan Agency for Medical Research and Development (AMED) [23bm1123026h0001], by grants from JST SPRING grant and supported by the Suzuki Kenzo Memorial Foundation for the Advancement of Medical Sciences, the Osaka Medical Research Foundation for Intractable Diseases, SENSHIN Medical Research Foundation, Japan Heart Foundation Research Grant on Dilated Cardiomyopathy, The NOVARTIS Foundation (Japan) for the Promotion of Science, Miyata Foundation Bounty for Pediatric Cardinovasucular Research, Mochida Memorial Foundation for medical and Pharmaceutical Research, and Kobayashi Foundation.

## Author Contributions

K.M., S.T., and O.T. conceptualized the project and designed the experiments. H.I., H.J., and H.H. performed CRISPRa/i experiments and gene expression analyses in cardiomyocytes. K.M., H.I. and H.J. performed ATAC-seq and ChIP-seq experiments. K.M., H.I., and H.J. performed RNA-FISH and 4C-seq analyses. Y.K. and O.T. generated and validated the CR9-deleted iPSC line and performed the iPSC-CM experiments. K.M., C.O., Y.M., Y.A., and O.T. generated and maintained the CR9 knockout and knock-in mouse lines. K.M., H.Y.F., and T.S. conducted phenylephrine stimulation studies and echocardiography. H.K., S.N., and Y.S. collected human myocardial samples. K. M., H. I., H.J., R.T., and O. T. engaged in extensive discussions regarding the structure and framing of the manuscript. K. M., H. I., H.J., and O. T. analyzed the data and wrote the manuscript with input from all authors. K.M., S.T., and O.T. secured funding and coordinated the collaborative efforts.

## Competing interests

The authors declare no competing interests.

## Notes

### Competing Interest Statement

The authors have declared no competing interest.

## References

1. Heidenreich PA, Bozkurt B, Aguilar D, Allen LA, Byun JJ, Colvin MM, Deswal A, Drazner MH, Dunlay SM, Evers LR, et al. 2022 AHA/ACC/HFSA Guideline for the Management of Heart Failure: A Report of the American College of Cardiology/American Heart Association Joint Committee on Clinical Practice Guidelines. Circulation. 2022;145:e895–e1032. doi: 10.1161/CIR.0000000000001063

2. Matsuoka K, Asano Y, Higo S, Tsukamoto O, Yan Y, Yamazaki S, Matsuzaki T, Kioka H, Kato H, Uno Y, et al. Noninvasive and quantitative live imaging reveals a potential stress-responsive enhancer in the failing heart. FASEB J. 2014;28:1870–1879. doi: 10.1096/fj.13-245522

3. Miyashita Y, Tsukamoto O, Matsuoka K, Kamikubo K, Kuramoto Y, Ying Fu H, Tsubota T, Hasuike H, Takayama T, Ito H, et al. The CR9 element is a novel mechanical load-responsive enhancer that regulates natriuretic peptide genes expression. FASEB J. 2021;35:e21495. doi: 10.1096/fj.202002111RR

4. Man JCK, van Duijvenboden K, Krijger PHL, Hooijkaas IB, van der Made I, de Gier-de Vries C, Wakker V, Creemers EE, de Laat W, Boukens BJ, et al. Genetic Dissection of a Super Enhancer Controlling the Nppa-Nppb Cluster in the Heart. Circ Res. 2021;128:115–129. doi: 10.1161/CIRCRESAHA.120.317045

5. Kuhn M, Voss M, Mitko D, Stypmann J, Schmid C, Kawaguchi N, Grabellus F, Baba HA. Left ventricular assist device support reverses altered cardiac expression and function of natriuretic peptides and receptors in end-stage heart failure. Cardiovasc Res. 2004;64:308–314. doi: 10.1016/j.cardiores.2004.07.004

6. Grosveld F, van Staalduinen J, Stadhouders R. Transcriptional Regulation by (Super)Enhancers: From Discovery to Mechanisms. Annu Rev Genomics Hum Genet. 2021;22:127–146. doi: 10.1146/annurev-genom-122220-093818

7. Schoenfelder S, Fraser P. Long-range enhancer-promoter contacts in gene expression control. Nature reviews Genetics. 2019;20:437–455. doi: 10.1038/s41576-019-0128-0

8. Blobel GA, Higgs DR, Mitchell JA, Notani D, Young RA. Testing the super-enhancer concept. Nature reviews Genetics. 2021;22:749–755. doi: 10.1038/s41576-021-00398-w

9. Hnisz D, Abraham BJ, Lee TI, Lau A, Saint-Andre V, Sigova AA, Hoke HA, Young RA. Super-enhancers in the control of cell identity and disease. Cell. 2013;155:934–947. doi: 10.1016/j.cell.2013.09.053

10. Whyte WA, Orlando DA, Hnisz D, Abraham BJ, Lin CY, Kagey MH, Rahl PB, Lee TI, Young RA. Master transcription factors and mediator establish super-enhancers at key cell identity genes. Cell. 2013;153:307–319. doi: 10.1016/j.cell.2013.03.035

11. Perlman BS, Burget N, Zhou Y, Schwartz GW, Petrovic J, Modrusan Z, Faryabi RB. Enhancer-promoter hubs organize transcriptional networks promoting oncogenesis and drug resistance. Nature communications. 2024;15:8070. doi: 10.1038/s41467-024-52375-6

12. Bender MA, Ragoczy T, Lee J, Byron R, Telling A, Dean A, Groudine M. The hypersensitive sites of the murine beta-globin locus control region act independently to affect nuclear localization and transcriptional elongation. Blood. 2012;119:3820–3827. doi: 10.1182/blood-2011-09-380485

13. Thomas HF, Kotova E, Jayaram S, Pilz A, Romeike M, Lackner A, Penz T, Bock C, Leeb M, Halbritter F, et al. Temporal dissection of an enhancer cluster reveals distinct temporal and functional contributions of individual elements. Mol Cell. 2021;81:969–982 e913. doi: 10.1016/j.molcel.2020.12.047

14. Hornblad A, Bastide S, Langenfeld K, Langa F, Spitz F. Dissection of the Fgf8 regulatory landscape by in vivo CRISPR-editing reveals extensive intra- and inter-enhancer redundancy. Nature communications. 2021;12:439. doi: 10.1038/s41467-020-20714-y

15. Shin HY, Willi M, HyunYoo K, Zeng X, Wang C, Metser G, Hennighausen L. Hierarchy within the mammary STAT5-driven Wap super-enhancer. Nat Genet. 2016;48:904–911. doi: 10.1038/ng.3606

16. Huang J, Li K, Cai W, Liu X, Zhang Y, Orkin SH, Xu J, Yuan GC. Dissecting super-enhancer hierarchy based on chromatin interactions. Nature communications. 2018;9:943. doi: 10.1038/s41467-018-03279-9

17. Hay D, Hughes JR, Babbs C, Davies JOJ, Graham BJ, Hanssen L, Kassouf MT, Marieke Oudelaar AM, Sharpe JA, Suciu MC, et al. Genetic dissection of the alpha-globin super-enhancer in vivo. Nat Genet. 2016;48:895–903. doi: 10.1038/ng.3605

18. Li K, Liu Y, Cao H, Zhang Y, Gu Z, Liu X, Yu A, Kaphle P, Dickerson KE, Ni M, et al. Interrogation of enhancer function by enhancer-targeting CRISPR epigenetic editing. Nature communications. 2020;11:485. doi: 10.1038/s41467-020-14362-5

19. Donovan LJ, Brewer CL, Bond SF, Laslavic AM, Pena Lopez A, Colman L, Jordan CE, Hansen LH, Gonzalez OC, Pujari A, et al. Aging and injury drive neuronal senescence in the dorsal root ganglia. Nat Neurosci. 2025;28:985–997. doi: 10.1038/s41593-025-01954-x

20. Alexander KA, Yu R, Skuli N, Coffey NJ, Nguyen S, Faunce CL, Huang H, Dardani IP, Good AL, Lim J, et al. Nuclear speckles regulate functional programs in cancer. Nat Cell Biol. 2025;27:322–335. doi: 10.1038/s41556-024-01570-0

21. Roadmap Epigenomics C, Kundaje A, Meuleman W, Ernst J, Bilenky M, Yen A, Heravi-Moussavi A, Kheradpour P, Zhang Z, Wang J, et al. Integrative analysis of 111 reference human epigenomes. Nature. 2015;518:317–330. doi: 10.1038/nature14248

22. Dijkstra JR, Opdam FJ, Boonyaratanakornkit J, Schonbrunner ER, Shahbazian M, Edsjo A, Hoefler G, Jung A, Kotsinas A, Gorgoulis VG, et al. Implementation of formalin-fixed, paraffin-embedded cell line pellets as high-quality process controls in quality assessment programs for KRAS mutation analysis. J Mol Diagn. 2012;14:187–191. doi: 10.1016/j.jmoldx.2012.01.002

23. Heist T, Fukaya T, Levine M. Large distances separate coregulated genes in living Drosophila embryos. Proc Natl Acad Sci U S A. 2019;116:15062–15067. doi: 10.1073/pnas.1908962116

24. Benabdallah NS, Williamson I, Illingworth RS, Kane L, Boyle S, Sengupta D, Grimes GR, Therizols P, Bickmore WA. Decreased Enhancer-Promoter Proximity Accompanying Enhancer Activation. Mol Cell. 2019;76:473–484 e477. doi: 10.1016/j.molcel.2019.07.038

25. Chen H, Levo M, Barinov L, Fujioka M, Jaynes JB, Gregor T. Dynamic interplay between enhancer-promoter topology and gene activity. Nat Genet. 2018;50:1296–1303. doi: 10.1038/s41588-018-0175-z

26. Lim B, Levine MS. Enhancer-promoter communication: hubs or loops? Curr Opin Genet Dev. 2021;67:5–9. doi: 10.1016/j.gde.2020.10.001

27. Du M, Stitzinger SH, Spille JH, Cho WK, Lee C, Hijaz M, Quintana A, Cisse, II. Direct observation of a condensate effect on super-enhancer controlled gene bursting. Cell. 2024;187:2595–2598. doi: 10.1016/j.cell.2024.04.001

28. Boija A, Klein IA, Sabari BR, Dall’Agnese A, Coffey EL, Zamudio AV, Li CH, Shrinivas K, Manteiga JC, Hannett NM, et al. Transcription Factors Activate Genes through the Phase-Separation Capacity of Their Activation Domains. Cell. 2018;175:1842–1855 e1816. doi: 10.1016/j.cell.2018.10.042

29. Sabari BR, Dall’Agnese A, Boija A, Klein IA, Coffey EL, Shrinivas K, Abraham BJ, Hannett NM, Zamudio AV, Manteiga JC, et al. Coactivator condensation at super-enhancers links phase separation and gene control. Science. 2018;361. doi: 10.1126/science.aar3958

30. Yoshimi K, Kunihiro Y, Kaneko T, Nagahora H, Voigt B, Mashimo T. ssODN-mediated knock-in with CRISPR-Cas for large genomic regions in zygotes. Nature communications. 2016;7:10431. doi: 10.1038/ncomms10431

31. Bulger M, Groudine M. Functional and mechanistic diversity of distal transcription enhancers. Cell. 2011;144:327–339. doi: 10.1016/j.cell.2011.01.024

32. Honnell V, Norrie JL, Patel AG, Ramirez C, Zhang J, Lai YH, Wan S, Dyer MA. Identification of a modular super-enhancer in murine retinal development. Nature communications. 2022;13:253. doi: 10.1038/s41467-021-27924-y

33. Gasperini M, Tome JM, Shendure J. Towards a comprehensive catalogue of validated and target-linked human enhancers. Nature reviews Genetics. 2020;21:292–310. doi: 10.1038/s41576-019-0209-0

34. Fulco CP, Nasser J, Jones TR, Munson G, Bergman DT, Subramanian V, Grossman SR, Anyoha R, Doughty BR, Patwardhan TA, et al. Activity-by-contact model of enhancer-promoter regulation from thousands of CRISPR perturbations. Nat Genet. 2019;51:1664–1669. doi: 10.1038/s41588-019-0538-0

35. Kassouf MT, Francis HS, Gosden M, Suciu MC, Downes DJ, Harrold C, Larke M, Oudelaar M, Cornell L, Blayney J, et al. The alpha-globin super-enhancer acts in an orientation-dependent manner. Nature communications. 2025;16:1033. doi: 10.1038/s41467-025-56380-1

36. Warren SA, Terada R, Briggs LE, Cole-Jeffrey CT, Chien WM, Seki T, Weinberg EO, Yang TP, Chin MT, Bungert J, et al. Differential role of Nkx2-5 in activation of the atrial natriuretic factor gene in the developing versus failing heart. Mol Cell Biol. 2011;31:4633–4645. doi: 10.1128/MCB.05940-11

37. Ostuni R, Piccolo V, Barozzi I, Polletti S, Termanini A, Bonifacio S, Curina A, Prosperini E, Ghisletti S, Natoli G. Latent enhancers activated by stimulation in differentiated cells. Cell. 2013;152:157–171. doi: 10.1016/j.cell.2012.12.018

38. Nord AS, Blow MJ, Attanasio C, Akiyama JA, Holt A, Hosseini R, Phouanenavong S, Plajzer-Frick I, Shoukry M, Afzal V, et al. Rapid and pervasive changes in genome-wide enhancer usage during mammalian development. Cell. 2013;155:1521–1531. doi: 10.1016/j.cell.2013.11.033

39. He A, Gu F, Hu Y, Ma Q, Ye LY, Akiyama JA, Visel A, Pennacchio LA, Pu WT. Dynamic GATA4 enhancers shape the chromatin landscape central to heart development and disease. Nature communications. 2014;5:4907. doi: 10.1038/ncomms5907

40. Core LJ, Martins AL, Danko CG, Waters CT, Siepel A, Lis JT. Analysis of nascent RNA identifies a unified architecture of initiation regions at mammalian promoters and enhancers. Nat Genet. 2014;46:1311–1320. doi: 10.1038/ng.3142

41. Mahat DB, Tippens ND, Martin-Rufino JD, Waterton SK, Fu J, Blatt SE, Sharp PA. Single-cell nascent RNA sequencing unveils coordinated global transcription. Nature. 2024;631:216–223. doi: 10.1038/s41586-024-07517-7

42. Hnisz D, Shrinivas K, Young RA, Chakraborty AK, Sharp PA. A Phase Separation Model for Transcriptional Control. Cell. 2017;169:13–23. doi: 10.1016/j.cell.2017.02.007

43. Fukaya T, Lim B, Levine M. Enhancer Control of Transcriptional Bursting. Cell. 2016;166:358–368. doi: 10.1016/j.cell.2016.05.025

44. Lim B, Heist T, Levine M, Fukaya T. Visualization of Transvection in Living Drosophila Embryos. Mol Cell. 2018;70:287–296 e286. doi: 10.1016/j.molcel.2018.02.029

45. Ma H, Tu LC, Naseri A, Chung YC, Grunwald D, Zhang S, Pederson T. CRISPR-Sirius: RNA scaffolds for signal amplification in genome imaging. Nat Methods. 2018;15:928–931. doi: 10.1038/s41592-018-0174-0

46. Gacita AM, Fullenkamp DE, Ohiri J, Pottinger T, Puckelwartz MJ, Nobrega MA, McNally EM. Genetic Variation in Enhancers Modifies Cardiomyopathy Gene Expression and Progression. Circulation. 2021;143:1302–1316. doi: 10.1161/CIRCULATIONAHA.120.050432

